# Multi-omics characterization of mesenchymal stem/stromal cells for the identification of putative critical quality attributes

**DOI:** 10.1101/2021.05.10.440010

**Authors:** Ty S. Maughon, Xunan Shen, Danning Huang, Adeola O Adebayo Michael, William A. Shockey, Seth H. Andrews, Jon M. McRae, Manu O Platt, Facundo M. Fernández, Arthur S. Edison, Steven L. Stice, Ross A. Marklein

**Author notes:** Equally contributing authors (Ty S. Maughon, Xunan Shen). **Corresponding Authors:** Dr. Ross Marklein and Dr. Steven Stice, Full Postal Address: 425 River Rd, Athens, GA 30602.

## Abstract

**Background:** Mesenchymal stromal cells (MSCs) have shown great promise in the field of regenerative medicine as many studies have shown that MSCs possess immunomodulatory function. Despite this promise, no MSC therapies have been granted licensure from the FDA. This lack of successful clinical translation is due in part to MSC heterogeneity and a lack of critical quality attributes (CQAs). While MSC Indoleamine 2,3-dioxygnease (IDO) activity has been shown to correlate with MSC function, multiple CQAs may be needed to better predict MSC function.

**Methods:** Three MSC lines (two bone marrow, one iPSC) were expanded to three passages. At the time of harvest for each passage, cell pellets were collected for nuclear magnetic resonance (NMR) and ultra-performance liquid chromatography mass spectrometry (UPLC-MS), and media was collected for cytokine profiling. Harvested cells were also cryopreserved for assessing function using T cell proliferation and IDO activity assays. Linear regression was performed on functional and multiomics data to reduce the number of important features, and partial least squares regression (PLSR) was used to obtain putative CQAs based on variable importance in projection (VIP) scores.

**Results:** Significant functional heterogeneity (in terms of T cell suppression and IDO activity) was observed between the three MSC lines, as well as donor-dependent differences based on passage. Omics characterization revealed distinct differences between cell lines using principal component analysis (PCA). Cell lines separated along principal component 1 based on tissue source (bone marrow vs. iPSC-derived) for NMR, MS, and cytokine profiles. PLSR modeling of important features predicts MSC functional capacity with NMR (R^2^=0.86), MS (R^2^=0.83), cytokines (R^2^=0.70), and a combination of all features (R^2^=0.88).

**Discussion:** The work described here provides a platform for identifying putative CQAs for predicting MSC functional capacity using PLSR modeling that could be used as release criteria and guide future manufacturing strategies for MSCs and other cell therapies.

## Introduction

Mesenchymal stem/stromal cells (MSCs) have been explored as a cell therapy in clinical trials due to their immunomodulatory properties.^1^ Although MSCs have shown great promise in preclinical studies for treatment of immune diseases, there have been challenges translating MSCs into approved therapies. This lack of translation can be attributed to MSC heterogeneity and no well-established critical quality attributes (CQAs i.e. limits or ranges of MSC biological properties) used to monitor MSC functional capacity among cell-lines or even within MSC cultures.^2–4^ T cell suppression is one of the most commonly used assays to assess MSC immunomodulatory capacity, but it is not standardized, has low throughput, and there is donor-donor variability among peripheral blood mononuclear cell (PBMC) responses to stimulation.^5^

The International Society for Cell and Gene Therapy (ISCT) has proposed several candidate protein properties to use for predicting MSC functional capacity.^2^ Indoleamine 2,3-dioxygenase (IDO) is one indicator of potency that has been proposed that plays a major role in the mechanisms by which MSCs modulate immune cells, such as T cells.^6^ Although IDO correlates with T cell suppression, there are several other mechanisms through which MSCs exert their potent immunomodulatory effects.^7–9^ Therefore, a matrix-based approach (*i.e.* a combination of CQAs) is likely needed to ensure a high quality MSC product.^2^ Chinnadurai *et al.* examined the relationship between various RNAs and secreted molecules with T cell proliferation and showed that several secreted factors and RNAs had strong correlations with T cell proliferation.^10^ Although these factors correlated with MSC functional capacity, measurement of these cellular properties require stimulation of sample MSC product acquired at the end of the manufacturing process and thus cannot be performed in-process.

MSCs secrete a broad repertoire of immunomodulatory cytokines that may have therapeutic potential; however, these immunomodulatory functions have not yet been characterized. Several studies have shown that MSC secreted cytokines can be modulated by physiologic conditions such as hypoxia, and pharmacological conditions such as targeted small molecule and growth factor conditioning.^11–13^ MSCs from different sources may differ in their immunomodulatory capacities, and the ability to assess cytokine secretion in MSCs using a non-destructive approach such as cytokine assays will allow for a standardized metric for MSC potency during manufacturing;^14^ cytokines that are secreted into the media can be measured without interference with cell growth and other conditions. Therefore, combination of specific cytokines secreted by MSCs, together with metrics of T cell suppression and IDO may enable identification of non-destructive, in-process CQAs for predicting potency.

Measures of cellular metabolism are also promising for assessing MSC quality due to the high abundance of metabolites in cells, and their importance in stem cell fate.^15^ Studies have shown that extended *in vitro* culture of MSCs shifts their metabolism from glycolysis towards oxidative phosphorylation (OXPHOS). MSCs in their native environment have a more glycolytic metabolism, and it has been shown that glycolytic MSCs have improved immunomodulatory effects *in vivo*.^15–17^ Therefore, assessing metabolite profiles in MSCs during expansion could be used as a predictor of their immunomodulatory capacity. Non-targeted metabolomics enables a detailed profiling of therapeutic cells, providing opportunities towards a more precise understanding of cellular therapeutic mechanisms.^18^ Nuclear magnetic resonance (NMR) spectroscopy and mass spectrometry (MS) are the two most commonly used techniques in metabolomics.^19^ NMR requires minimal sample preparation, making it highly reproducible. NMR can also provide information in assigning metabolites based on chemical shifts and J-coupling patterns. As an analytical platform, MS also has certain advantages. Its higher sensitivity enables the detection of low abundance metabolites that are below the NMR detection thresholds,^20^ whereas its high resolution greatly reduces spectral overlap. When coupled with separation techniques such as gas chromatography (GC) or ultra-performance liquid chromatography (UPLC), spectral complexity is greatly reduced, and metabolic chemical properties can be revealed.^21–23^ The combination of NMR and UPLC-MS metabolic profiling provides an even more in-depth measurement of MSC cellular metabolism, potentially leading to the discovery of CQAs. To date, NMR- and MS-based metabolic profiling was used to characterize cellular metabolism, leading to the discovery of biomarkers or pathways different between cell lines or cellular responses of treatment.^24–26^ However, to the best of our knowledge, no studies have been reported where NMR- and/or UPLC-MS-based metabolomics are used to establish CQAs associated with MSC immunomodulatory potency.

In this study, we used a multi-omics approach to identify MSC metabolites and cytokine levels during cell manufacturing that could serve as predictors of MSC immunomodulation and as potential potency assays. These metabolites and cytokines are referred to in this work as putative CQAs. Intracellular metabolites and secreted cytokines from three MSC lines at multiple passages were studied to determine their correlation with MSC immunosuppressive capacity post thaw. NMR, UPLC-MS, and cytokine data sets were also merged and filtered to identify putative CQAs. Using partial least squares regression (PLSR) and variable importance projection (VIP) scores, we identified a multi-omic panel of candidate cytokines and metabolites that could be used to predict MSC functional capacity and inform future manufacturing strategies.

## Methods

### Cell Culture

Two bone marrow-derived MSC lines (RoosterBio, Frederick MD) (lot #0071 and #0182, which the manufacturer has both research and clinical-grade lots available), and one induced pluripotent stem cell derived MSC cell-line (Cellular Dynamics International, Madison WI) (Lot #0003, also prequalified) were used and referred here as BM71, BM182, and iMSC, respectively. MSCs were thawed and allowed to recover for 24 hours in complete medium (MSC-GM) (Alpha-Minimum Essential Medium (Gibco), 10% fetal bovine serum (Hyclone), 2mM L-glutamine, 50 U/mL penicillin, 50 μg/mL streptomycin (Gibco)) before being seeded at 500 cells/cm^2^. Cells were expanded in 10 150 mm plates with a negative, media-only control plate and 14 T175 flasks for expansion. After reaching approximately 80% confluency, cells in 150 mm dishes were washed with PBS three times. Cells were then scraped and collected in 80:20 methanol:water solution and stored at −80°C until further analysis. Cells grown in T175 flasks for expansion were harvested using 0.05% trypsin (Gibco) and counted using a Cellometer K2 cell counter (Nexcelom, Lawrence MA). Cells were either cryopreserved for functional assays or reseeded in dishes/flasks for continued expansion (3 total passages for each cell-line, 9 total experimental groups). Population doubling level (PDL) for each cell-line/passage was determined using formula 1:

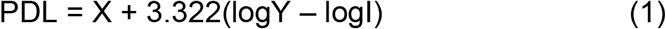

Where X = initial PDL, I = initial cell seeding number and Y = final number of cells.

### T Cell Suppression Assay

MSCs from each cell-line/passage were thawed and allowed to recover for 48 hours with a media change at 24 hours. MSCs were then seeded at three densities (10,000, 5,000 and 2,000 cells/well) in a 96 well plate and cultured for 24 hours. PBMCs (AllCells, Alameda CA) were thawed in RPMI media (RPMI, 20% FBS, 2mM L-glutamine, 50 U/mL penicillin, 50 μg/mL streptomycin) and cultured overnight at 37°C and 5% CO_2_. Prior to co-culture, PBMCs were labeled with CFSE (Supplemental Table 1, Biolegend, San Diego CA) according to the manufacturer’s protocol, and 100,000 PBMCs were added to each well at final MSC:PBMC ratios of 1:10, 1:20, or 1:50 as well as control wells containing only PBMCs. Following PBMC addition, stimulating anti-CD3/CD28 Dynabeads (Thermo Fisher Scientific, Waltham MA) were added at 100,000 beads per well to the appropriate wells (positive controls and all MSC groups). MSCs and PBMCs were co-cultured for 72 hours at 37°C, 5% CO_2_.

Following co-culture, PBMCs were collected and stained using APC/Fire anti-CD4 and APC anti-CD8 (Supplemental Table 1) (Biolegend, San Diego CA). PBMCs were first washed and stained with Zombie Yellow (Supplemental Table 1) (Biolegend, San Diego CA) viability dye and blocked using 2% FBS. PBMCs were then washed again and stained for CD4 and CD8 in the dark at room temperature. Following staining, the antibodies were blocked using 2% FBS and washed. Cells were then fixed with 4% PFA for 30 minutes at 4°C. Cells were then washed and re-suspended in PBS containing 2% FBS. Cells were stored overnight at 4°C in the dark until flow analysis.

### Flow Cytometry

All flow cytometry experiments were performed using a CytoFLEX S (Beckman Coulter, Hialeah FL) with 20,000 events collected per sample. All flow cytometry data were analyzed using FlowJo (Treestar, Inc., Ashland OR). Briefly, cell debris, doublets, and Dynabeads were gated out using scatter principles. Then, single stained controls were used for compensation, and fluorescence minus one controls were used in order to determine positive populations (Supplemental Figure 1).

### IDO Activity Assay

MSCs from each cell-line/passage experimental group were thawed and cultured for 48 hours with a media change at 24 hours. MSCs were then seeded at a density of 40,000 cells/cm^2^ in a 96 well plate in MSC-GM. After 24 hours, the medium was replaced in each well with MSC-GM containing 10 ng/mL interferon gamma (IFN-γ) (Life Technologies). After an additional 24 hours, conditioned media was collected and frozen at −20°C, and cells were fixed with 4% PFA. Media was thawed and 100 μL was transferred into a 96 well plate. Trichloroacetic acid was used to precipitate excess protein. 75 μL of the supernatant was collected and transferred to a separate 96 well plate. Ehrlich’s Reagent was then added to each well to detect L-kynurenine levels using a SpectraMax iD5 (Molecular Devices) plate reader. Levels of L-kynurenine were determined using a standard curve. To normalize L-kynurenine values to cell numbers for each experimental group, we performed automated image analysis to quantify cell nuclei in the wells from which conditioned medium was collected. Following fixation, MSCs were washed with PBS twice, and stained with Hoechst. Cells were then imaged on a Cytation 5 high content imaging system (Biotek, Winooski VT) and cell counts determined using CellProfiler^27^ to normalize the amount of L-kynurenine per cell.

### Metabolomics Sample Preparation

MSC samples were thawed at 4°C and vortexed three times for 1 min. Then samples were centrifuged at 14,000 x g for 5 min at 4°C. For each sample, 30% (300 μl) and 60% (600 μl) of the supernatant were transferred to pre-labeled Eppendorf tubes for LC-MS and NMR spectroscopy, respectively. For each PDL, 67μl of each sample was pooled together to generate 2 internal PDL quality control (QC) samples. The remaining 33 μl supernatant of each sample was pooled together to generate 2 internal overall QC samples. In each PDL, the extraction blank sample was added using extraction solvent (methanol: water 80:20). Samples were then evaporated in a Speedvac for 6 hours and stored in −80°C until future analysis. NMR samples with QC controls and 2 buffer blank samples were used for data acquisition. Samples were all randomized, with total 49 samples in each cell line. The LC-MS sample randomization was identical to NMR.

### NMR

The NMR buffer solution was prepared by dissolving 928.6 mg of anhydrous NaH_2_PO_4_ and 320.9 mg of Na_2_HPO_4_ in 80 ml D_2_O (Cambridge Isotope Laboratory) in a volumetric flask. Sodium trimethylsilylpropanesulfonate (DSS) was used as a chemical shift and concentration reference standard by adding 333.3 μl of 1.0 M DSS-D_6_ (Cambridge Isotope Laboratory) stock solution to the buffer for a final DSS concentration of 1/3 mM. The pH was adjusted to 7.4 (uncorrected for isotope effects) and was brought to a volume of 100 ml with D_2_O and mixed well. The pH was rechecked, and the buffer stored at 4°C until use.

The NMR samples were reconstituted in 80 μl of the NMR buffer and vortexed thoroughly. Sixty μl of each sample was transferred to racks of 96 1.7-mm NMR tubes for data acquisition using a SamplePro Tube robotic system (Bruker Biospin, Billerica, MA, USA). Samples were run on a Bruker NEO 800 MHz NMR spectrometer equipped with a 1.7-mm cryoprobe and Bruker SampleJet cooled to 5.6°C. One dimensional nuclear Overhauser enhancement spectroscopy with water suppression (1D-NOESY PR) was collected on all samples. The spectra were processed using NMRPipe^28^ and and in-house MATLAB metabolomics toolbox (https://github.com/artedison/Edison_Lab_Shared_Metabolomics_UGA). The spectra were aligned using Correlation Optimized Wrapping (COW) algorithm^29^ and normalized with a Probabilistic quotient normalization (PQN)^30^ algorithm. The non-overlapped peaks were manually binned, and the area under curve was calculated for each non-overlapped feature.

Two-dimensional ^1^H-^13^C Heteronuclear Single Quantum Coherence (HSQC) and ^1^H-^13^C HSQC-Total Correlated Spectroscopy (HSQC-TOCSY) spectra on internal pooled samples were collected for metabolite identification. The spectra were processed using NMRPipe, and the in-house MATLAB metabolomics toolbox. The 2D NMR spectra were matched to a metabolite database using the COLMARm.^31^ The metabolites were assigned a confidence level ranging from 1 to 5 according to published criteria.^32^

After metabolite identification, peaks were further normalized by a normalization factor calculated as the proton numbers in functional group corresponding to a given peak. A total of 28 metabolites were obtained from binning of the whole spectra. The unknown binned features were then used to perform correlation analysis. The features with correlation coefficient values greater or equal to 0.8 were then grouped together as tentative unknown metabolites. A total of 29 tentative unknown features were extracted from 100 features (Supplemental Figure 2).

### UPLC-MS

Cell extract samples were resuspended in 50 μL methanol/water (80:20 *v/v).* A sample blank was prepared with 50 μL of methanol/water solution (80:20 *v/v*), and a pooled quality control (QC) sample was created by mixing a 10 μL aliquot of each cell extract sample. Both the sample blank and the pooled sample were processed with the same procedure, and were analyzed together with the cell extract samples. All samples were run in randomized order on consecutive days. QC samples were analyzed every 10 runs to assess UPLC-MS system stability and correct time-dependent batch effects with a QC-based regression curve.

UPLC-MS analyses were performed using a Vanquish Horizon UPLC (Thermo Fisher Scientific, Inc., Waltham, MA) system coupled to a Orbitrap ID-X Tribrid mass spectrometer (Thermo Fisher Scientific, Inc., Waltham, MA). Hydrophilic interaction (HILIC) chromatography was performed with a Waters ACQUITY UPLC BEH HILIC, 2.1 × 75 mm, 1.7 μm particle column. Mobile phase A was water/acetonitrile (95:5 v/v), 10 mM ammonium acetate, and 0.05% ammonium hydroxide. Mobile phase B was acetonitrile with 0.05% ammonium hydroxide. Chromatographic gradients can be found in Supplemental Information (Supplemental Table 2). The column temperature was 55 °C, while samples were maintained at 5 °C in the autosampler. Two μL of each sample were injected, and the mass spectrometer was operated in positive ion mode. For metabolite identification purposes, data-dependent acquisition (DDA) experiments were used to collect MS/MS spectra at stepped normalized collision energy (NCE) of 10, 30, and 50.

### Cytokine Profiling

Conditioned media from each cell-line/passage (at the time of cell harvest) were collected and prepared for cytokine profiling using the Human Premixed Multi-Analyte Magnetic Luminex Assay (R&D Systems, Minnneapolis MN) (Supplemental Table 3). Cytokine profiling was performed following a standard protocol. Briefly, 50 μL standard or media sample was added to a 96-well plate, followed by 50 μL diluted microparticle cocktail and incubated for 2-hours at room temperature on an orbital microplate shaker. Using a magnetic device attached to the bottom of the microplate, each well was washed with 50 μl of wash buffer. Then 50 μL diluted biotin-antibody cocktail was added to wells and incubated for 1-hour at room temperature on an orbital microplate shaker. After washing, 50 μL of streptavidin-PE was added to each well and incubated for 30 min. 100μL of wash buffer was added to wells to resuspend microparticles and samples were then loaded into a BioPlex 200 Luminex system (Bio-Rad) and analyzed. The assay measures the intensity of PE emission, which is correlated to cytokine concentration standards.

### Data Analysis and Statistical Methods

Principal component analysis (PCA) was conducted using JMP (version 15). All suppression assays, linear regression models, and IDO assays were analyzed using Prism (GraphPad, San Diego CA). Comparison of T cell proliferation to the positive control, comparison of T cell proliferation among all groups at the 1:10 dilution, and IDO activity were analyzed using one-way ANOVA with Tukey’s post-hoc test. Comparisons of T cell suppression against passage 1 and against the 1:10 dilution were analyzed using a two-way ANOVA with Tukey’s post hoc test.

NMR and MS features were mapped to CD4^+^ and CD8^+^ proliferation rates using simple linear regression. The top 20 metabolites of each dataset with the lowest p-values (p<0.05) were selected for downstream analysis. Selected 20 NMR metabolites, 20 MS metabolites and all cytokine data were used to train a partial least squares regression (PLSR) model. The response was the PCA loading score generated by using principal component 1 of all five functional measures termed, composite functional score. The predictors with variable importance in projection (VIP) scores greater or equal to 1 were selected as potential CQAs.

## Results

### Cell-line and passage-dependent differences in T cell suppression

Population doubling level was used to assess MSC age and growth characteristics for each cell-line throughout expansion (Supplemental Table 4). Population doubling level, doublings per day, and log growth of cell expansions were recorded (Figure 1B-D). Following expansion, MSC cell-lines at three different passages were co-cultured with anti-CD3/CD28 stimulated PBMCs in order to assess their immunosuppressive capacity with two PBMC donors. MSCs were cultured at three different MSC:PBMC ratios as well as positive (no MSCs with stimulation) and negative (no MSCs, no stimulation) controls. After 3 days, CD4^+^ and CD8^+^ T cell proliferation was assessed based on CFSE dilution (Figure 2A-F). A dose response was observed when increasing the MSC:PBMC ratio on the ability of the iMSC and BM182 cells to suppress CD4^+^ and CD8^+^ proliferation (p<0.05). CD4^+^ and CD8^+^ T cell proliferation decreased at the 1:50 dilution compared to the 1:10 dilution (p<0.05) in BM71 except for P3. The 1:10 MSC:PBMC ratio had the greatest variation in function across all cell-lines and passages and was used for comparison and predictive function. All passages of the iMSCs and BM182 lines suppressed CD4^+^ and CD8^+^ T cell proliferation when compared to the positive control (p<0.05) (Figure 2A,C,D,F). BM71 increased CD4^+^ proliferation at all passages and increased CD8^+^ proliferation at P1 (p<0.05) and had no significant differences at P2 and P3 when compared to the positive control (Figure 2B,E). For BM182, CD4^+^ and CD8^+^ T cell proliferation (p<0.05) was suppressed at P3 compared to earlier passages and had similar suppression as the iMSCs (Figure 2G). These studies were repeated using PBMCs harvested from a second donor (Supplemental Figure 2). Similarly, there was a MSC dose dependent response, as the 1:10 MSC:PBMC ratio had significantly lower (p<0.05) CD4^+^ and CD8^+^ proliferation when compared with 1:20 and 1:50 ratios (Supplemental Figure 2 A-F). Although BM71 did suppress both CD4^+^ (except P1) and CD8^+^ T cells when compared to the positive control (p<0.05), they possessed different immunomodulatory capacities than the iMSCs and BM182 (CD4^+^ and CD8^+^ p<0.05) at the 1:10 ratio (Supplemental Figure 2G,H). Again, iMSCs consistently suppressed both CD4^+^ and CD8^+^ proliferation across all passages. BM182 P3 had significantly less CD4^+^ proliferation compared to P1 (p<0.05) and P2 (p<0.05). BM182 P3 also had significantly less CD8^+^ proliferation compared to P1 (p<0.05), was less than P2 although not technically significant (p=0.0597) (Supplemental Figure 2G,H).

**Figure 1.**
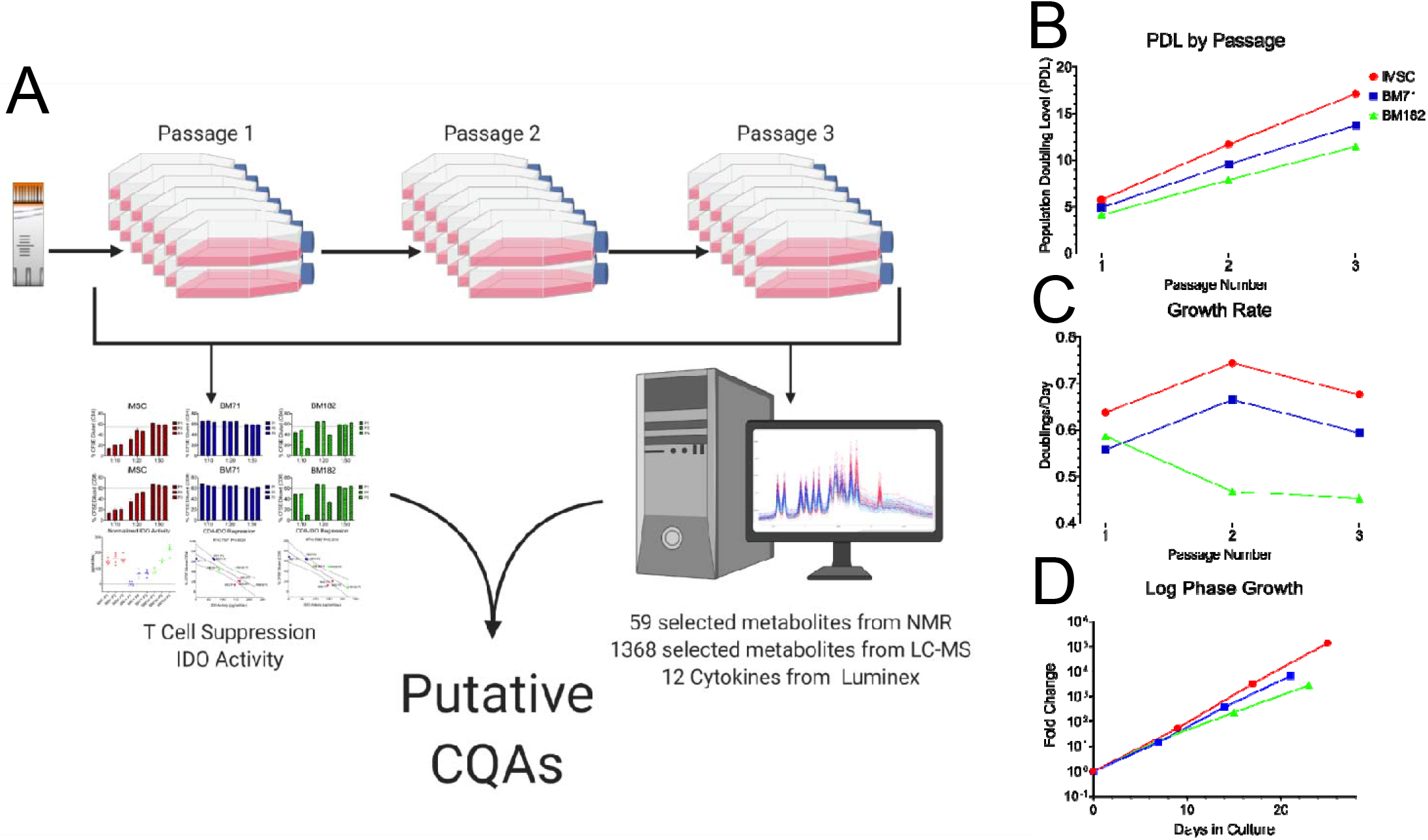
MSC expansion workflow and growth characteristics. (A) MSC characterization workflow from expanded cells to discover putative CQAs. (B) Population doubling level changes of three MSC lines over three passages. (C) MSC growth rate in doublings per day over three passages. (D) Log phase growth characteristics of each MSC line over the number of days in culture.

**Figure 2.**
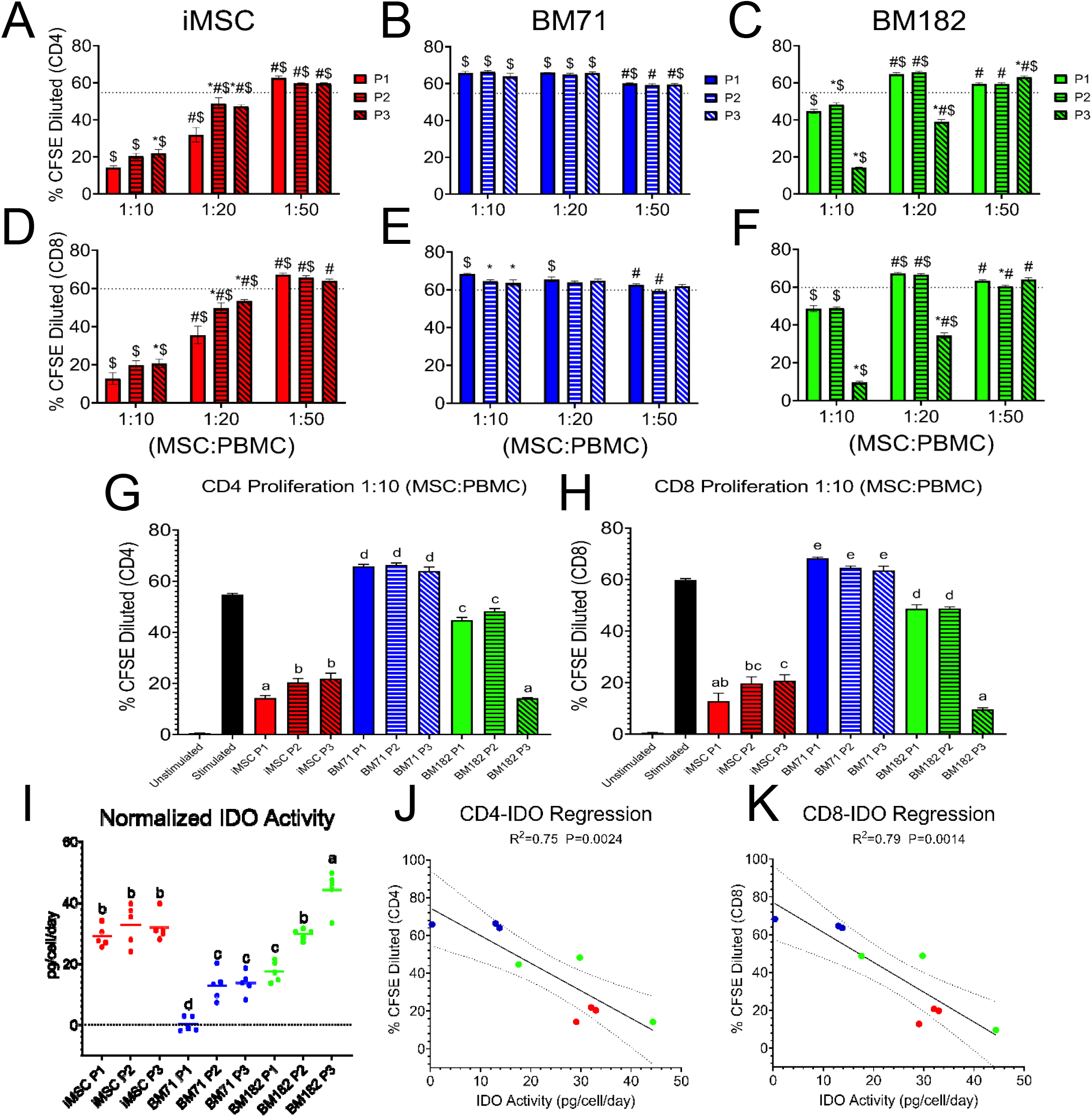
MSC functional capacity characterized by T-cell proliferation and IDO activity. MSCs were co-cultured with stimulated PBMCs at three different MSC:PBMC ratios (1:10, 1:20, 1:50). (A-C) CD4^+^ and (D-F) CD8^+^ T cell proliferation was assessed at each passage and ratio by % CFSE dilution. A 2-way ANOVA was used in order to determine if there was a significant difference from P1 within a ratio (*), a significant difference from the 1:10 ratio within a passage (#), or a significant difference from the stimulated control (dotted line)($)(P<0.05). (G) CD4^+^ and (H) CD8 T cell proliferation comparing all cell lines and passages at the 1:10 ratio, and (I) IDO activity measured by L-kynurenine levels normalized to cell number and days in culture were analyzed using a one-way ANOVA with Tukey’s post hoc test to determine significance (P<0.05). Linear regression of the relationship between IDO activity and (J) CD4^+^ and (K) CD8^+^ T cell proliferation.

### MSC IDO activity correlates with T cell proliferation

IDO activity for each cell-line/passage group was quantified as another functional readout of MSC immunomodulatory capacity (Figure 2I). Similar to the T cell suppression results, iMSCs displayed high IDO activity (indicated by high levels of L-kynurenine) for all passages and BM71 had the lowest IDO activity for all cell-lines at each passage. IDO activity of BM182 increased from P1 to P2 (p<0.05) with P3 displaying the highest levels of L-kynurenine (P<0.05) compared to all other cell-lines. Linear regression was performed to determine the relationship between IDO activity and CD4^+^/CD8^+^ T cell proliferation (Figure 2J, K). IDO activity correlated with both CD4^+^ and CD8^+^ T cell proliferation (R^2^=0.75 and R^2^=0.79, respectively) for the 1:10 MSC:PBMC ratio. The second PBMC donor was consistent with the first PBMC donor, IDO activity and MSC suppression of CD4^+^/CD8^+^ T cells (R^2^=0.85 and R^2^=0.81, respectively) were positively correlated (*i.e* higher IDO activity means higher T cell suppression) (Supplemental Figure 3).

### NMR-based identification of MSC metabolic signatures

For each cell-line/passage, the intracellular products were analyzed using NMR metabolomics profiling. The heat map of all detected metabolites shows that the overall metabolite signature was cell line specific with some passage differences observed within cell-lines (Figure 3A). As an unbiased way of looking at the metabolic signature of MSCs, we used unsupervised PCA to examine differences in the studied cell lines. Unsupervised PCA revealed distinct metabolic differences between the cell lines (Figure 3B). Different tissue donor sources clearly separated along PC1 with iMSC samples grouped together on the positive side of PC1 and the both BM cell lines on the negative side. iMSCs were significantly different (p<0.05) from BM71 and BM182 along PC1, but there were no observable differences between BM71 and BM182 (Figure 3C). BM182 separated from BM71 along PC2 at P3 (p<0.05), which is the passage at which BM182 showed significant functional improvement (Figure 3D).

**Figure 3.**
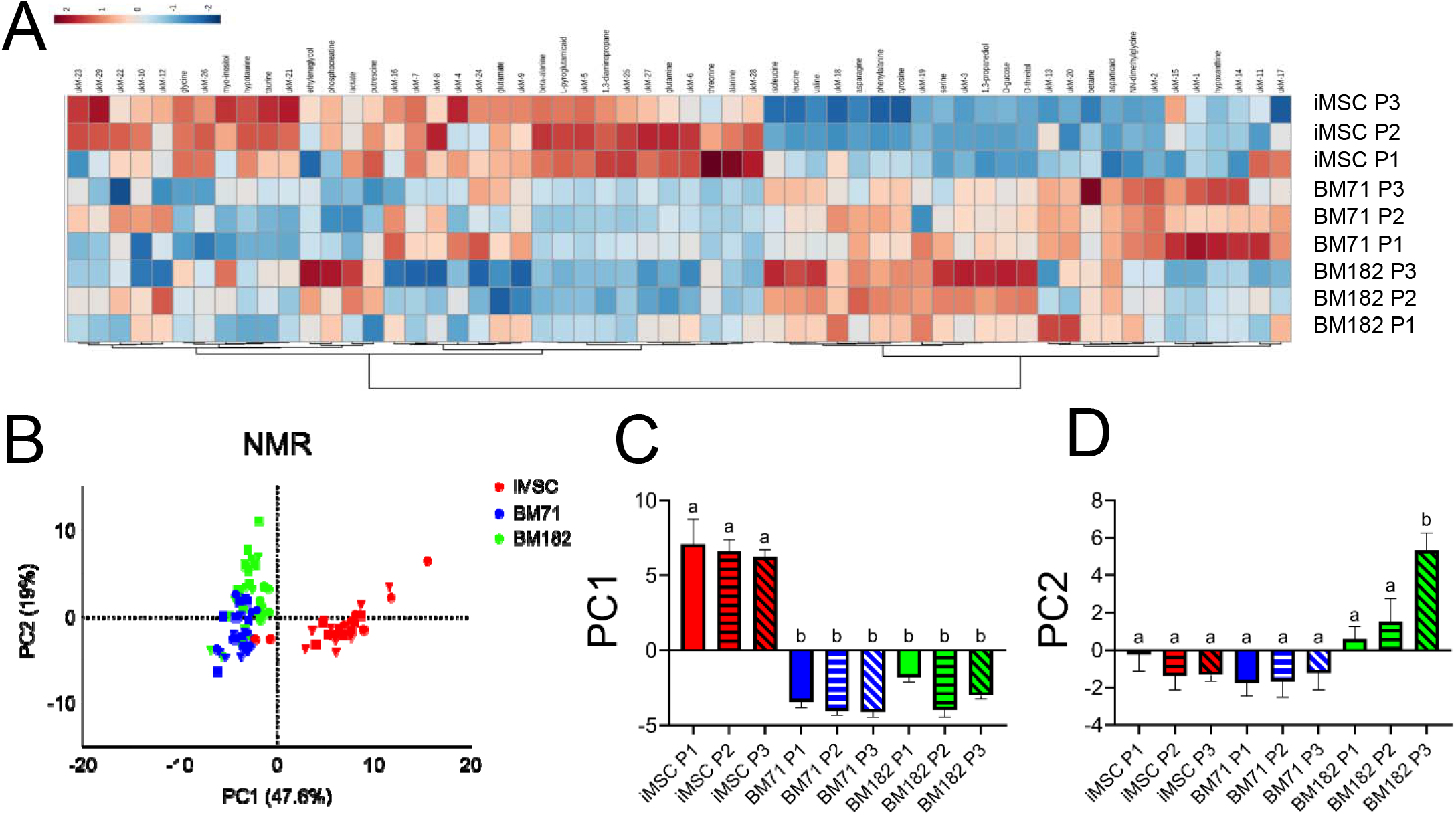
MSC metabolic profile from NMR analysis. (A) Heat map of all 58 metabolites using Euclidian distance measure and ward clustering method. (B) Unsupervised PCA of all metabolites for each cell line at three passages (P1=circle, P2=triangle, P3=square). (C) One-way ANOVA comparison of average PC1 value. (D) One-way ANOVA comparison of average PC2 value.

### UPLC-MS-based identification of MSC metabolic signatures

Additional cell pellet samples collected at the time of cell harvest for each cell line/passage were analyzed using UPLC-MS. As with NMR metabolic profiling results, the heat map for the 1368 metabolite features showed clear differences in the metabolic profile between BM and iMSC cell lines, and between different passages within a given cell line (Figure 5A). Similar to what was observed in NMR, PCA of the 1368 UPLC-MS metabolites displayed a clear separation of all cell lines (Figure 5B). iMSCs separated from both BM71 and BM182 (p<0.05) along the PC1 axis, with no differences between BM71 and BM182 (Figure 5C). All cell lines were significantly different (p<0.05) from one another along PC2 (Figure 5D). There were no metabolomic profile differences between passages within a cell line.

**Figure 4.**
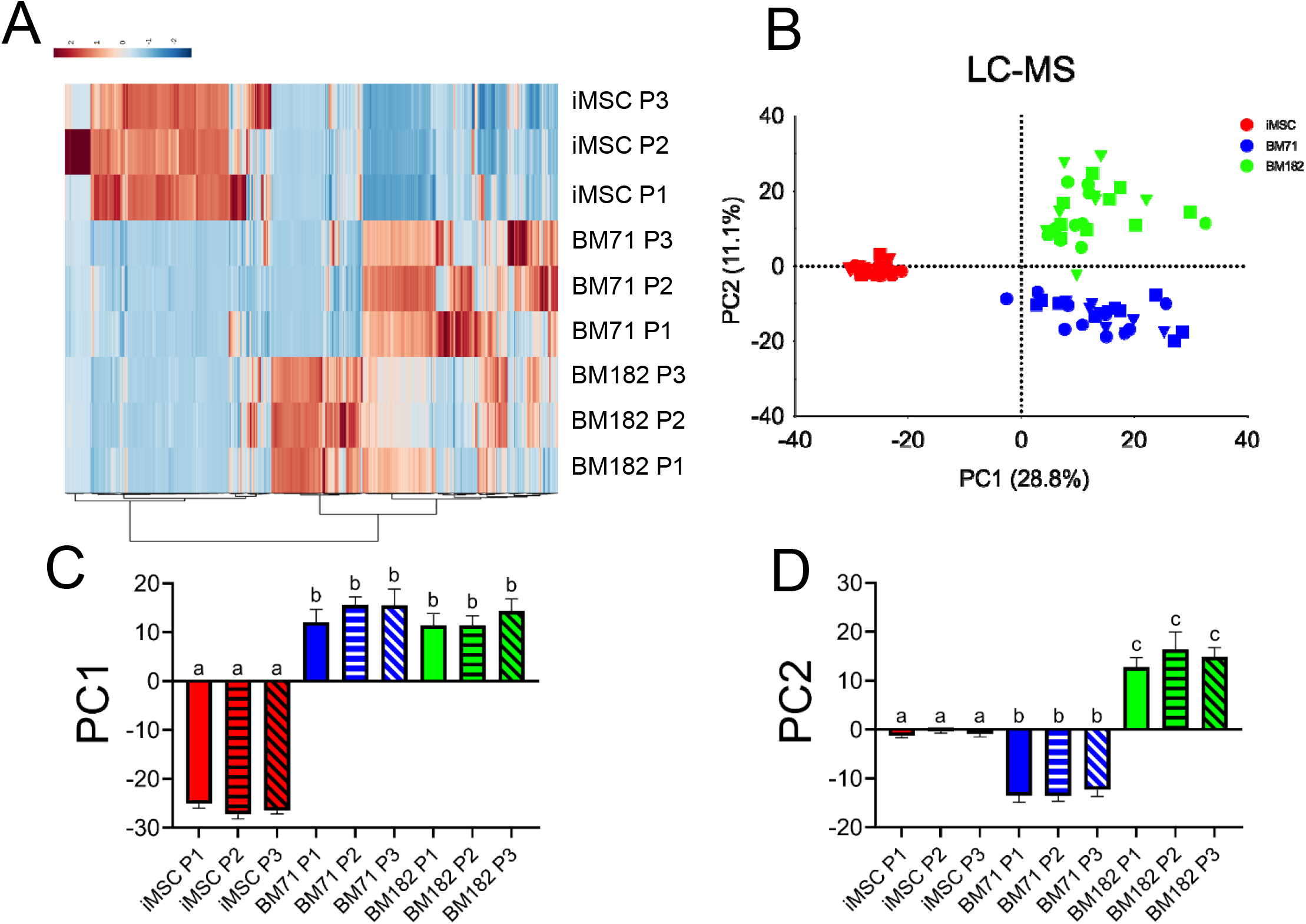
MSC metabolic profile from MS analysis. (A) Heat map of 1386 metabolites using Euclidian distance measure and ward clustering method. (B) Unsupervised PCA of all metabolites for each cell line at three passages (P1=circle, P2=triangle, P3=square). (C) One-way ANOVA comparison of average PC1 value. (D) One-way ANOVA comparison of average PC2 value.

**Figure 5.**
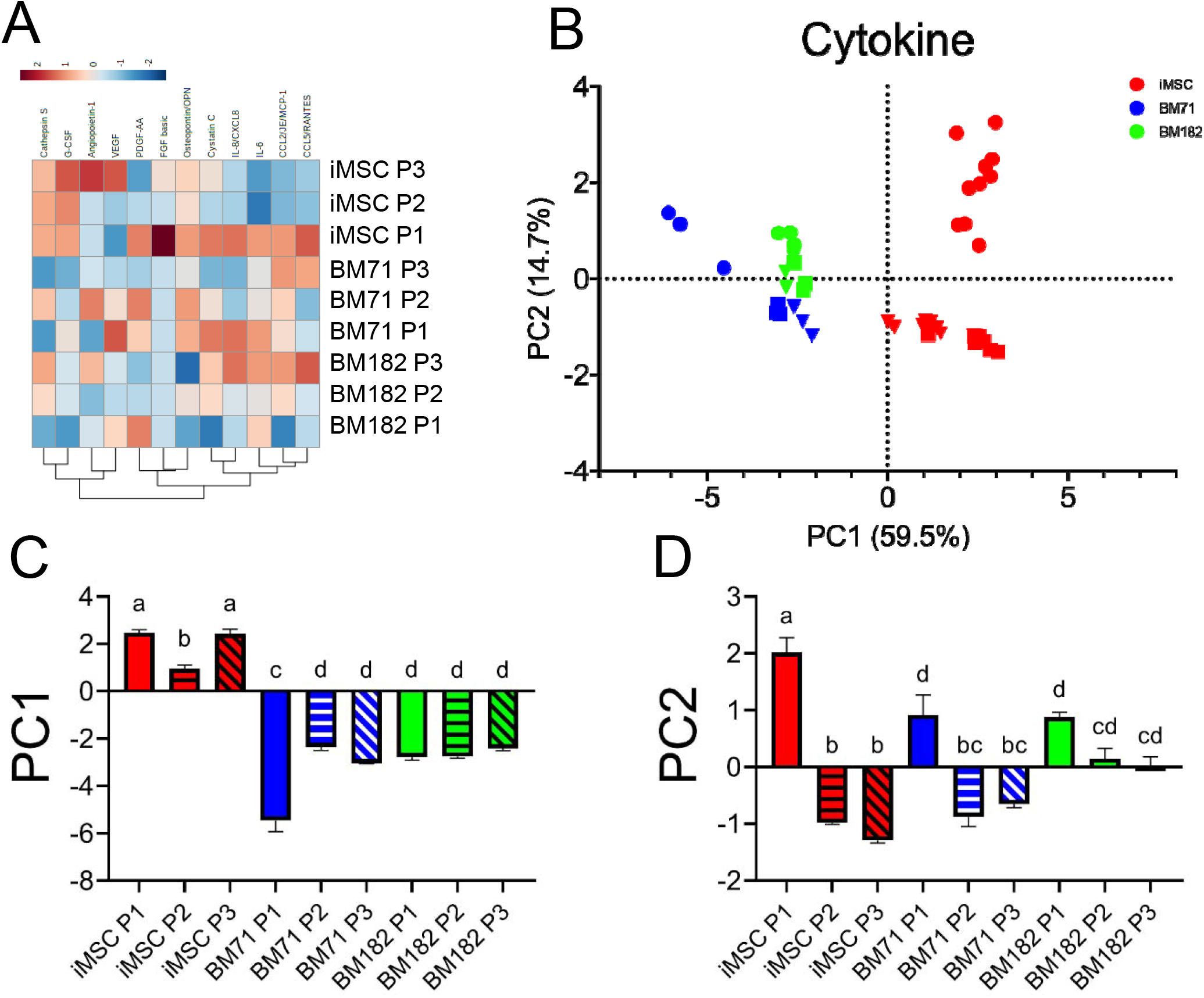
MSC cytokine profile. (A) Heat map of all 12 measured cytokines using Euclidian distance measure and ward clustering method. (B) Unsupervised PCA of all cytokines for each cell line at three passages (P1=circle, P2=triangle, P3=square). (C) One-way ANOVA comparison of average PC1 value. (D) One-way ANOVA comparison of average PC2 value.

### Secreted cytokine profile of MSC conditioned media

Conditioned media collected at the time of MSC harvest for each cell-line/passage was analyzed using Multi-Analyte Magnetic Luminex assay. The heatmap revealed variable cytokine expression from all cell lines (Figure 5A). Cytokines with similar functions clustered together, for example, CCL5 and CCL2, both chemotactic cytokines for T-cells are clustered. Similarly, VEGF and angiopoietin, angiogenic growth factors, are clustered on the heat map. IL-8, IL-6 and cystatin C clustered closely with the chemotactic cytokines (Figure 5A). To examine cytokine signature difference among the cell lines using an unbiased approach, we performed PCA. The scores plot showed a distinct separation of cytokine profiles among cell-lines (Figure 5B). The cytokine profiles of the iMSCs grouped together at PC1 on the positive side, while the two BM cell lines were clustered on the negative side (p<0.05) (Figure 5C). Differences were also observed within the iMSC line (P2 *vs.* P1/P3) and BM71 line (P1 *vs*. P2/P3)(p<0.05) (Figure 5C). The first passage of iMSCs and BM71 were significantly different (p<0.05) from the later passages within their respective cell-line along PC2 (Fig 5D). BM182 showed no significant differences along PC2 (Fig 5D).

### Regression of identified metabolites to determine putative CQAs for predictive function

Metabolites identified from NMR and MS were regressed with CD4^+^ and CD8^+^ T cell proliferation (Figure 6). Metabolites with strong positive or negative correlations (R^2^>0.50) were then used in order to reduce the data set and ordered based on the strongest correlation. Regression with PBMC Donor 1 revealed strong correlations with multiple NMR metabolites. Myo-inositol had the highest R^2^ values for both CD4^+^ (R^2^=0.84) and CD8^+^ (R^2^=0.84) T cell proliferation as followed by unknown NMR metabolite 2 (ukNMR-2) (R^2^=0.82 and R^2^=0.77, respectively) (Figure 6A-C). To confirm these results, NMR metabolites were then regressed with PBMC Donor 2. Myo-inositol and ukNMR-2 correlated (R^2^=0.54 and R^2^=0.74, respectively) with CD4^+^ proliferation (Supplemental Figure 5). Regression with CD8^+^ proliferation showed very similar results to PBMC Donor 1 with myo-inositol and ukNMR-2 having strong correlations (R^2^=0.81 and R^2^=0.88, respectively). Linear Regression with UPLC-MS metabolites also revealed multiple metabolites that correlated with function. From PBMC donor 1, the top two correlated metabolites for CD4^+^ and CD8^+^ proliferation were unknown MS metabolite 1349 (ukM-1349) (R^2^=0.88 and R^2^=0.85, respectively) and phosphatidylcholine, PC(O-38:4) (R^2^=0.81 and R^2^=0.88, respectively) (Figure 6E-H). ukM-1349 and PC(O-38:4) both correlated with PBMC donor 2 CD4^+^ (R^2^=0.64 and R^2^=0.55, respectively) and CD8^+^ (R^2^=0.87 and R^2^=0.83, respectively) T cell proliferation as well (Supplemental Figure 5).

**Figure 6.**
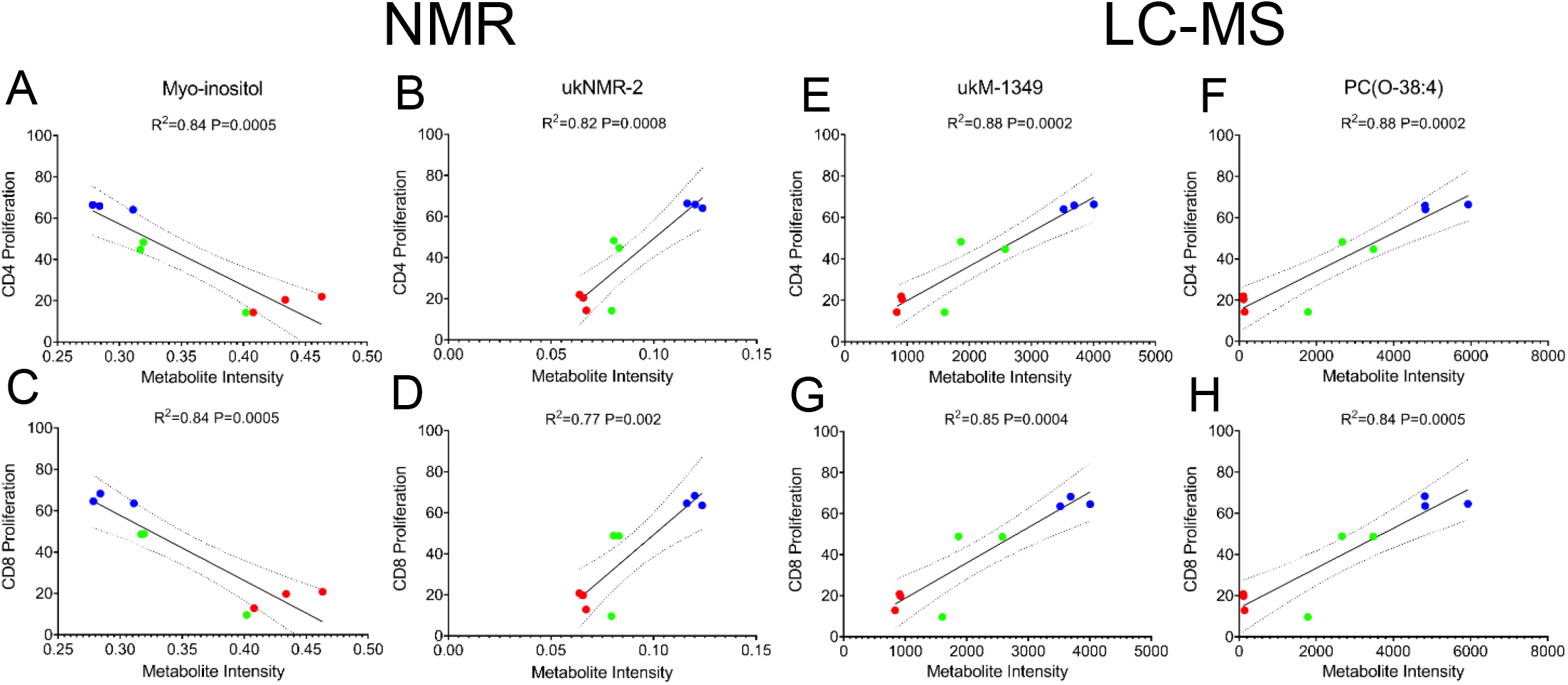
Linear regression analysis of the top correlated metabolites. (A,C) Myo-inositol and (B,D) ukNMR-2 had the highest R^2^ values for CD4^+^ and CD8^+^ T cell proliferation from NMR metabolites. (E,G) ukM-1349 and (F,H) PC(O-38:4) had the highest R^2^ values for CD4^+^ and CD8^+^ T cell proliferation from MS metabolites. The top 20 metabolites with the highest R^2^ values were then chosen for further modeling to discover putative CQAs. Red=iMSCs, Blue=BM71, Green=BM182

### PLSR modeling of putative CQAs enables prediction of MSC functional capacity

A composite functional score was created in order to assess each cell-line’s overall functional capacity due to varying levels of T cell activation from different PBMC donors (Figure 7A). All five functional outputs were plotted using PCA, and PC1 was then used as the composite functional score which accounted for 92.1% of the variance (Figure 7B,C). The top 20 correlated metabolites from linear regression analyses were then used to train the PLSR model. Features with VIP scores greater than 1 were selected: 6 NMR putative CQAs (Figure 7D), 7 MS putative CQAs (Figure 7E), 4 cytokine putative CQAs (Figure 7F), and 10 total putative CQAs when combining all data sets (Figure 7G) from each model respectively. PLSR was then retrained on each omics data set. NMR had the greatest R^2^ value (R^2^=0.86) followed by MS (R^2^=0.83) and cytokines (R^2^=0.70) (Figure 7H-J). Combining these data sets showed high predictability (R^2^=0.88), but not distinguishably higher than NMR or MS alone.

**Figure 7.**
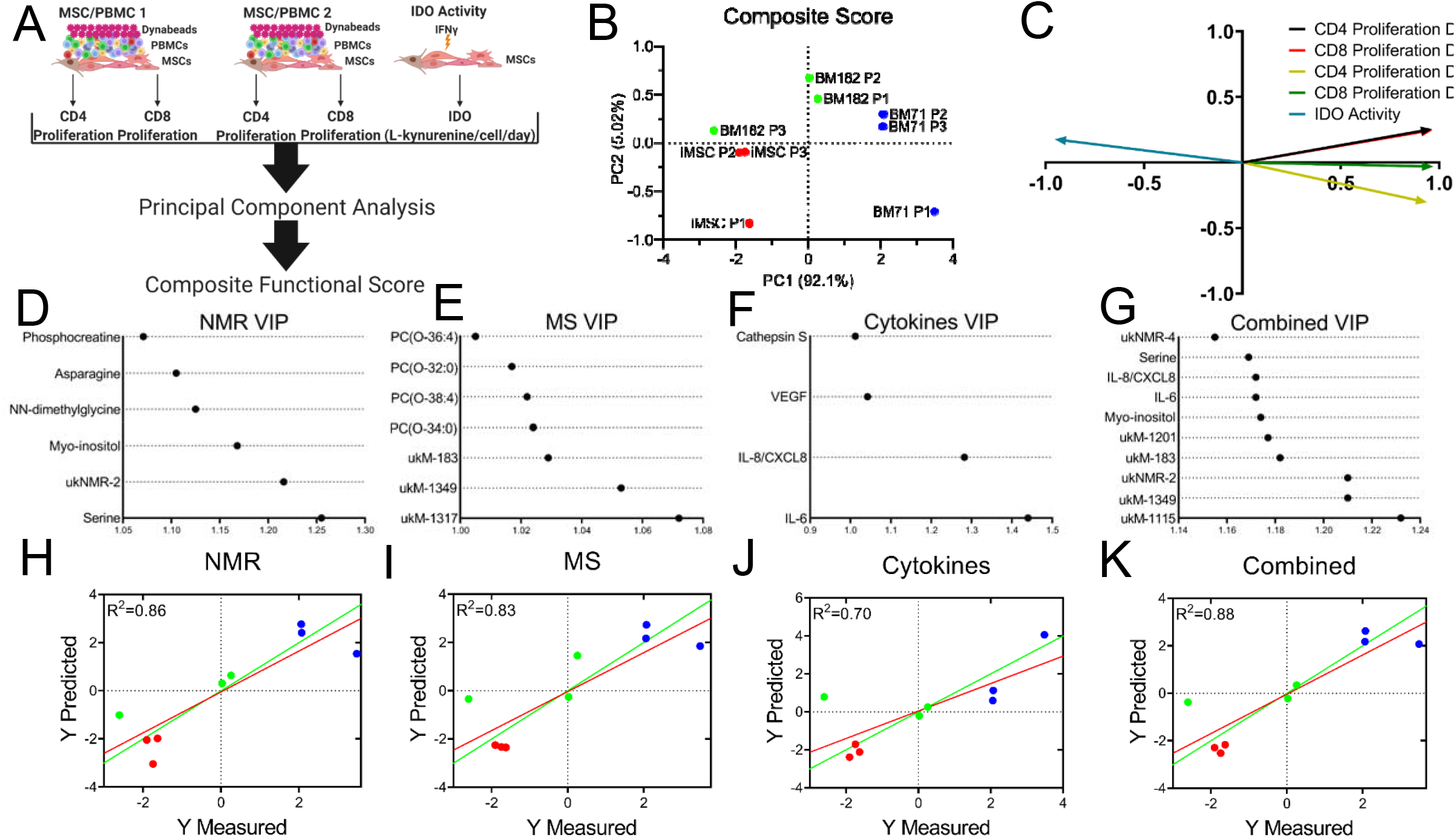
PLSR modeling of important features elucidates candidate CQAs. (A-C) CD4^+^ and CD8^+^ T cell proliferation from two donors and IDO activity (5 total functional outputs) were analyzed using PCA. PC1 accounted for 92.1% of the variance and was then taken and used as a composite functional score for further predictive modeling. VIP scores were assigned to the top (D) NM (E) MS, (F) cytokines, and (G) a combination of all features based on PLSR modeling (H-K). The VIP molecules chosen are considered as putative CQAs.

## Discussion

MSC therapies are currently being investigated due to their immunomodulatory properties, but translation to the clinic has been a challenge because of the heterogeneity of MSCs and a lack of CQAs that predict potency.^4,33^ Combinatorial, matrix-based approaches may help overcome these challenges by encompassing more than one functional metric since MSCs modulate the immune system using different mechanisms.^10^ In this study, we combined comprehensive metabolomics profiling with non-destructive cytokine profiling to determine several putative CQAs for assessing MSC immunosuppressive capacity. Using a composite functional score, these putative CQAs were strongly correlated with MSC functional capacity. The approach outlined in this study takes large data sets that sample distinct cellular functions and identifies important features for MSC potency prediction.

Cellular metabolism is a key regulator of MSC fate and immunomodulatory potency.^15^ Using an NMR based metabolomics approach, we found intracellular metabolites that positively and negatively correlated with MSC functional capacity. Amino acids such as serine and asparagine were significant based on a PLSR model with VIP selection. Serine has been shown to control the self-renewal of epidermal stem cells (EpdSCs). When extracellular serine is limited, EpdSCs activate de novo serine synthesis and promotes epidermal differentiation. Conversely, blocking serine synthesis facilitates malignant progression.^34^ Asparagine has been shown to regulate stem cell proliferation.^35^ Additionally, depletion of asparagine in cell culture medium results in diminished cell growth.^36^ NN-dimethylglycine, a derivative of the amino acid glycine, also positively correlated with MSC function. It is also a byproduct of the metabolism of choline. To our knowledge, there are no studies that demonstrate a relationship between NN-dimethylglycine and stem cell biology and thus further investigation is warranted. The role of myo-inositol is not known in cells, however myo-inositol is an important growth-promoting factor of mammalian cells, and possibly acts as an osmolyte.^37,38^ Myo-inositol constitutes a component of membrane phospholipids and mediates osmoregulation.^39^ All the identified metabolites play essential roles in cell growth and facilitate cell proliferation although their role in MSC immunomodulatory capacity needs to be further investigated.

MS-based metabolic profiling revealed 7 putative CQAs that were associated with MSC immunomodulatory potency. Four out of the 7 selected UPLC-MS CQAs were annotated as phosphatidylcholines (PCs), including PC(O-36:4), PC(O-38:4), and PC(O-32:0), and PC(O-34:0). PC constitutes a major portion of the cell membrane and play an important role in cellular reprogramming and signaling.^40^ The intermediate of PC synthesis or hydrolysis, lysophosphatidylcholines (LPCs), have been reported as markers for discriminating different MSC sources and could be related to differences in MSC differentiation capacity and immunomodulatory properties.^26^ The annotated PCs were observed to negatively correlate with MSC immunomodulatory capacity. Lower levels of PCs were detected in iMSCs at all passages and BM 182 at passage 3, which were the MSC groups that demonstrated higher immunosuppressive capacity. Furthermore, we discovered the 4 identified PCs group into 2 pairs (PC(O-38:4)/PC(O-36:4) and PC(O-34:0)/ PC(O-32:0)) sharing the same unsaturation degree and a difference of 2 carbons in their fatty acid chain composition, suggesting an underlying, yet unexplained, connection of those PC in their biosynthesis pathway. Increases in unsaturation levels of polyunsaturated PCs may alter membrane fluidity and hence can contribute to changes in MSC morphology, which has also been shown to predict MSC function.^41,42,43^ However, to date, the biological role of PCs in stem cell metabolism at the fatty acid chain level is still poorly understood.

MSCs have also been shown to have beneficial effects by secreting proteins to modulate cell behavior in regenerative medicine and health applications.^44–46^ For this study, cytokines that span multiple MSC functions of angiogenesis, tissue repair, and recruitment of immune cells were quantified.^47^ The amount of each secreted cytokine/growth factor was investigated in a non-destructive manner to enable monitoring of temporal changes due to cell secretion, media changes during expansion and passaging, as well as cell uptake through autocrine or paracrine signaling. Cytokine profiling in MSCs has been described in the literature as a metric of MSC functionality, with certain secreted cytokines upregulated in MSCs, and other cytokines unchanged or reduced, and secreted in multiple cell lines and donors.^10,48^ By investigating levels of cytokines in MSC culture media in conjunction with metabolites and T cell suppression, we have developed an assay matrix for predicting MSC potency for immunomodulation. Besides serving as potential effectors of MSC immunomodulatory function, the cytokines profiled in this study have also been shown to directly impact MSC behavior.^49,50^ A potential function is priming of MSCs, where MSCs are conditioned with specific cytokines to increase their immunomodulatory properties.^10^ Priming may be efficacious in instances where there is low MSC survival potential ex vivo or differences in sources and donor reduces effectiveness of MSC for use in regenerative and immunomodulatory applications. MSC priming using our in-process cytokines and culture conditions can be used to develop substrates or engineered tissue for regenerative medicine.^51^ Also, non-destructive monitoring of secreted factors in spent media could potentially serve as CQAs for large scale MSC manufacturing (in bioreactors, for example). MSC-secreted cytokines can regulate MSC function both in an autocrine and paracrine manner. For example, the binding of secreted IL-8 to its receptor CXCR1 or CXCR2 can activate intracellular PI3K, MAPK, Akt phosphorylation and initiate functions of cell survival, angiogenesis and cell migration.^52^ In contrast, targeted ablation of secreted cytokines such as IL-6, either by gene silencing or inhibition, led to reduced MSC proliferation and reduced capacity of MSCs to suppress T cell proliferation.^53^ Ultimately, studying this crosstalk between MSCs and secreted cytokine may be a relevant aspect of MSC expansion for manufacturing.

It is well documented that MSC functional heterogeneity is derived from differences in donor/tissue source, MSC doubling level, and manufacturing conditions.^33,54^ Being able to understand and predict these differences is a challenge that needs to be addressed in order to advance MSC therapies. Knowing what to measure can help screen for high potency MSCs and assess when these MSCs begin to lose potency due to senescence.^42,55,56^ Our multi-omics approach elucidated several correlative features that, when used in combination, were indicative of MSC potency and were able to predict an increase in potency in the BM182 line. IL-6 and IL-8 are both inflammatory cytokines that recruit immune cells such as T cells, neutrophils, and macrophages. They have also been shown to inhibit T cell apoptosis and regulatory T cell differentiation.^57–59^ Higher levels of these cytokines were secreted by less potent lines (BM71). Although the role of PCs in MSCs on immunomodulatory capacity is largely unknown, studies have shown that oxidized phospholipids, such as PCs, play a role in preventing the activation of T cells and dendritic cells as well as mediate apoptosis.^60–63^ Apoptosis of MSCs by cytotoxic T cells has been shown to play an important role in their immunomodulatory capacity^64^. Myo-inositol and serine’s role in MSC immune suppression has not been investigated to this point, but soluble myo-inositol has been shown to be effective in treating autoimmune diseases such as thyroiditis and hypothyroidism.^65^ Further investigation into the pathways involving these metabolites will improve our understanding of how MSCs modulate the immune system.

Effective prediction of MSC potency is an important factor in manufacturing high quality therapies. Predicting MSC functional capacity has been a major challenge because MSCs can exert their immunomodulatory effects through a number of different mechanisms and immune cells. Therefore, using a combination of cell metabolites and secreted cytokines can help better predict MSC potency and set specific CQAs to aid in process by design cell manufacturing systems.^2,4,10,66^ Our study used a combination of metabolites and cytokines in order to better predict MSC potency. These are easy to target and measure for manufacturing and understanding how these metabolites affect cell function can help refine and improve the manufacturing process. This approach of CQA discovery and understanding can also be translated into other cell therapies such as chimeric antigen receptor T-cell (CAR-T), iPSCs, neural stem cells, and even MSC-derived extracellular vesicles. Moving forward, interrogating pathways that involve these metabolites will be important for assay development and better understanding the relationship of MSC metabolism with immunomodulation. Non-destructive, in-process monitoring of MSC metabolism using conditioned medium will also enable on-demand, precise control of MSC manufacturing. Finally, this discovery platform can be used to establish MSC CQAs for other therapeutic applications involving differentiation (e.g. osteo-, adipo-, and chondrogenesis) and tissue engineering, cancer treatment, and angiogenesis.

## Abbreviations

MSC: Mesenchymal stem/stromal cells
CQA: critical quality attributes
PBMC: peripheral blood mononuclear cell
ISCT: The International Society for Cell and Gene Therapy
IDO: indoleamine 2,3-dioxygenase
OXPHOS: oxidative phosphorylation
NMR: nuclear magnetic resonance
MS: mass spectrometry
GC: gas chromatography
UPLC: ultra-performance liquid chromatography
PLSR: partial least squares regression
VIP: variable importance projection
IFN-γ: interferon gamma
1D-NOESY PR: one dimensional nuclear Overhauser enhancement spectroscopy with water suppression
COW: correlation optimized wrapping
PQN: probabilistic quotient normalization
HSQC: heteronuclear single quantum coherence
HSQC-TOCSY: HSQC-total correlated spectroscopy
HILIC: hydrophilic interaction chromatography
DDA: data-dependent acquisition
NCE: normalized collision energy
PCA: principal component analysis
PCs: phosphatidylcholines
ukNMR-n: unknown NMR metabolite n
ukM-n: unknown MS metabolite n
EpdSCs: epidermal stem cells
CAR-T: chimeric antigen receptor T-cell

## Acknowledgements

The authors thank Julie Nelson and the University of Georgia CTEGD Cytometry Shared Resource Laboratory for providing equipment and expertise and the Center for Undergraduate Research (CURO) at the University of Georgia for supporting Jon McRae III. NSF EEC-1648035 funded this work. All illustrations were made using biorender.com. FMF also acknowledges support from NSF MRI CHE-1726528. ASE and SS acknowledge support from the Georgia Research Alliance.

## Dataset

Detailed experimental NMR methods, as well as all raw and processed data are available on the Metabolomics Workbench (http://www.metabolomicsworkbench.org/), DataTrack ID: 2519.

## Author Contributions

Conception and design of the study: TM, XS, DH, WAS, MOP, FF, AE, RM, SS. Acquisition of data: TM, XS, DH, WAS, AOAM, JM, SA. Analysis and interpretation of data: TM, XS, DH, AOAM, MOP, FF, AE, RM, SS. Drafting or revising the manuscript: TM, XS, DH, AOAM, MP, FF, AE, RM, SS. All authors have revised and approved the final article.

---

**Supplemental Table 1.** PBMC co-culture assay antibodies and flow cytometer information.

| Laser | EM Filter | Marker | Color / Format | Host / Target | Isotype | Clone | Company | Catalog |
| --- | --- | --- | --- | --- | --- | --- | --- | --- |
| 638 | 780/30 | CD4 | APC/Fire | Mouse anti-Human | IgG1 κ | RPA-T4 | BioLegend | 67-0047-T025 |
| 638 | 660/20 | CD8 | APC | Mouse anti-Human | IgG1 | CN9V1 | BioLegend | orb248718 |
| 488 | 525/40 | CFSE | CFSE | N/A | N/A | N/A | BioLegend | 423801 |
| 405 | 610/20 | Zombie Yellow | Zombie Yellow | N/A anti-All Species | N/A | N/A | BioLegend | 423104 |

**Supplemental Figure 1.**
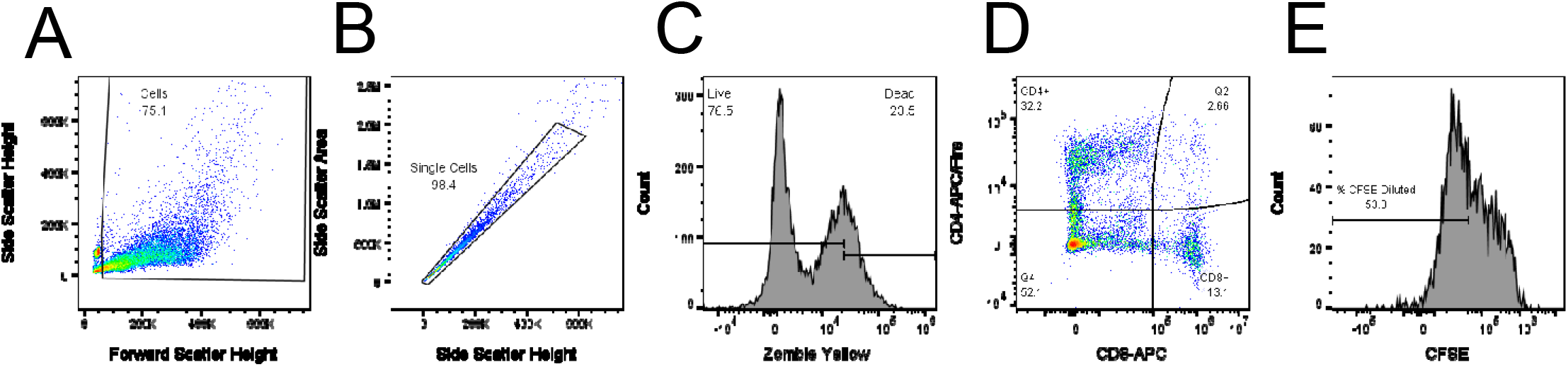
FlowJo gating strategy. Cellular debris and Dynabeads (A) were first removed followed by cell doublets (B). Using FMO controls, live cells (C) were then gated and used to determine CD4^+^ and CD8^+^ T cell populations (D). Negative control PBMCs (No stimulation, no MSCs) were then used for the CFSE dilution gate (E).

**Supplemental Figure 2.**
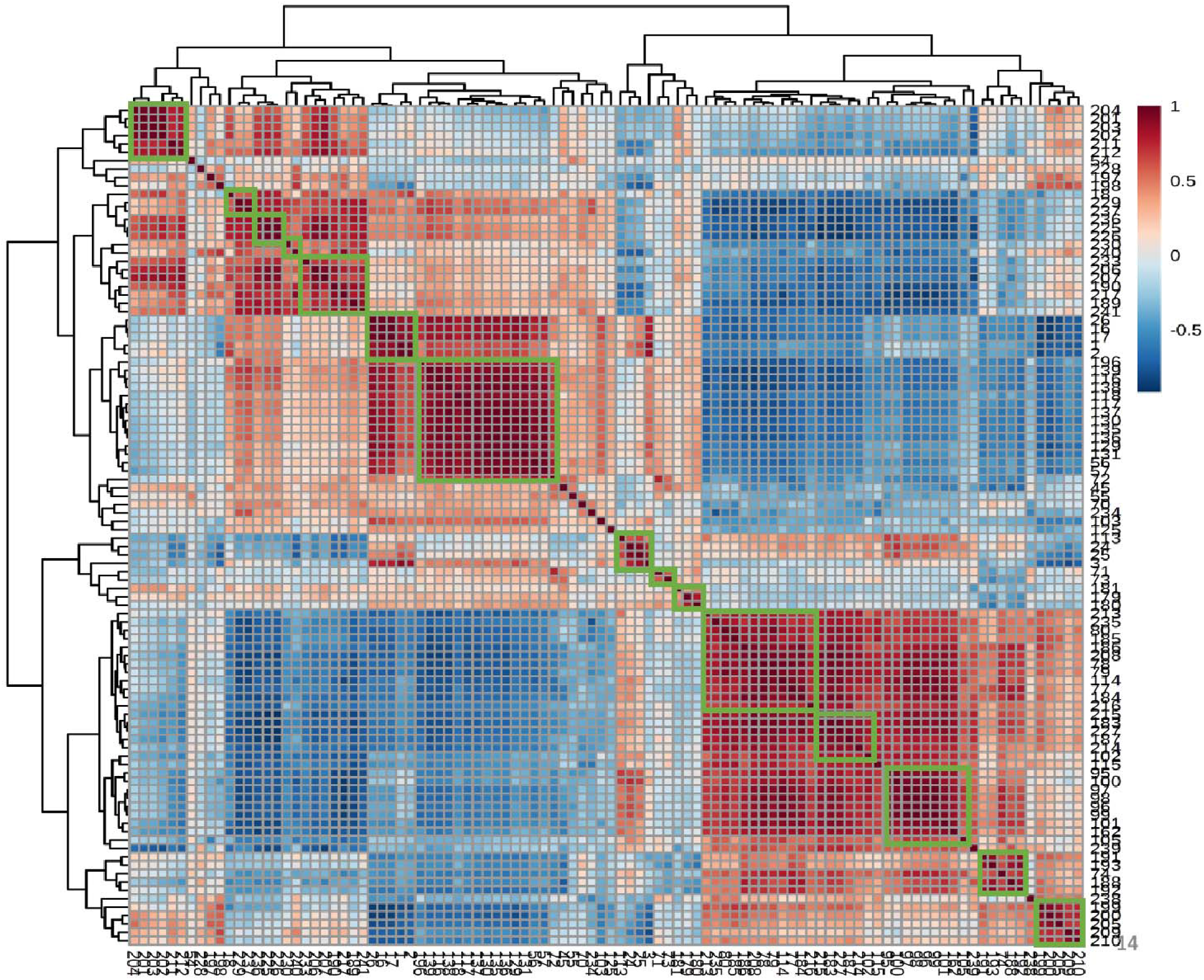
The correlation map of 100 unknown NMR features. We assume that correlation coefficient values greater or equal to 0.8 indicating those features are from same metabolite and grouped together as Indicated by green box in the figure. The features with correlation coefficient value less than 0.8, we considered them as individual metabolites. Total 100 unknown features were grouped to 29 unknown metabolites.

**Supplemental Table 2.**
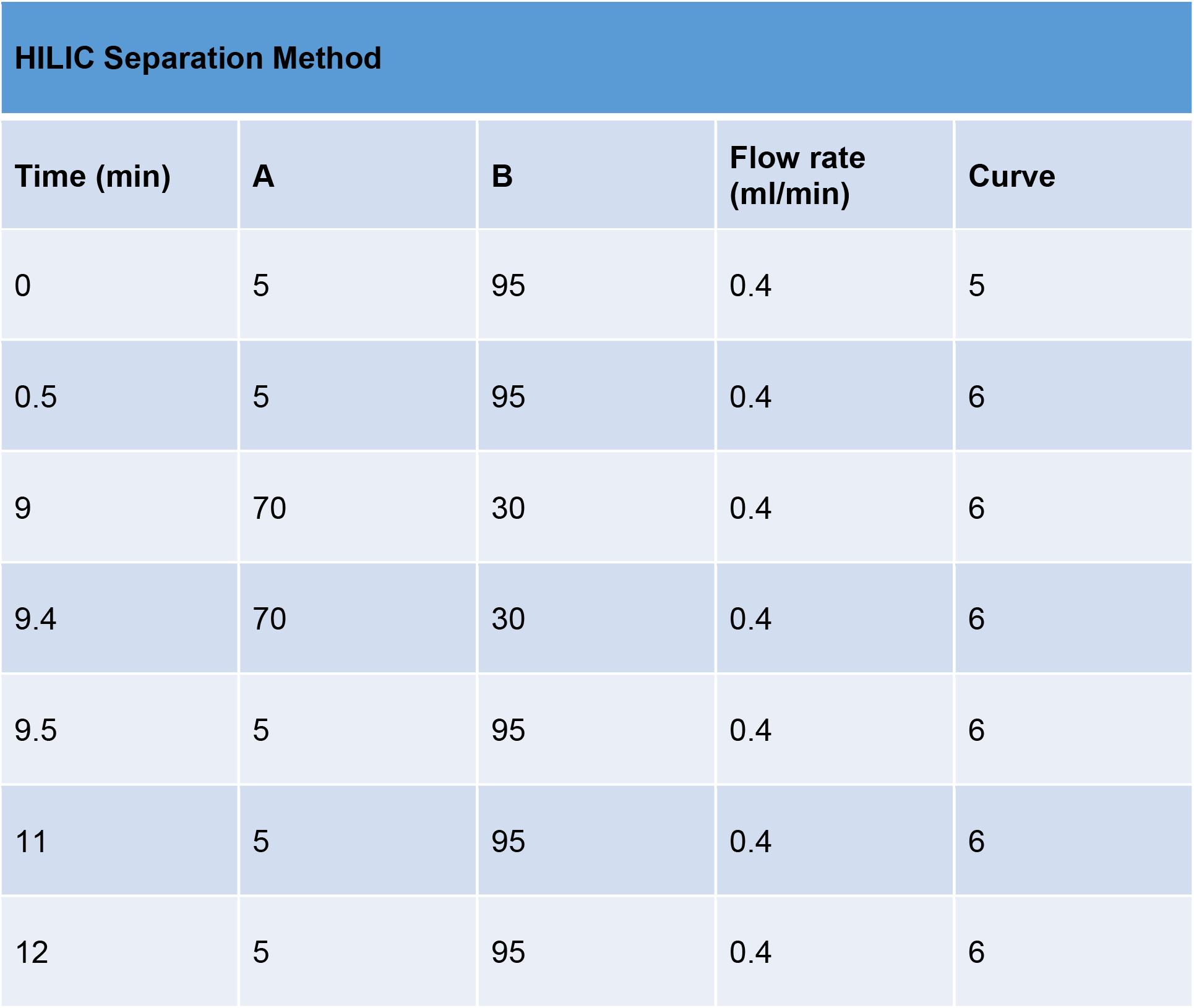
Methods information. Chromatographic gradient for HILIC method: mobile phase A was water/acetonitrile (95:5 v/v), with 10 mM ammonium acetate and 0.05% ammonium hydroxide, and B was acetonitrile with 0.05% ammonium hydroxide. Both ionization modes utilized identical mobile phases and identical chromatographic gradients.

HILIC Separation Method
| Time (min) | A | B | Flow rate (ml/min) | Curve |
| --- | --- | --- | --- | --- |
| 0 | 5 | 95 | 0.4 | 5 |
| 0.5 | 5 | 95 | 0.4 | 6 |
| 9 | 70 | 30 | 0.4 | 6 |
| 9.4 | 70 | 30 | 0.4 | 6 |
| 9.5 | 5 | 95 | 0.4 | 6 |
| 11 | 5 | 95 | 0.4 | 6 |
| 12 | 5 | 95 | 0.4 | 6 |

**Supplemental Table 3.** List of all cytokines measured and their corresponding function.

| Analyte | Function |
| --- | --- |
| Angiopoietin-1 | Promote angiogenesis and vascular growth |
| CCL2/MCP-1 | Recruits immune cells to sites of inflammation |
| Cystatin C | Inhibitor of cysteine cathepsin proteases |
| IL-8/CXCL8 | Stimulates neutrophil migration and phagocytosis |
| PDGF-AA | Cell growth, differentiation and survival |
| Cathepsin S | Proteolysis, antigen presentation, extracellular functions |
| CCL5/RANTES | Recruits immune cells to sites of inflammation |
| FGF basic | Numerous, cell growth and development, embryonic development |
| IL-6 | Numerous, pro-inflammatory signaling |
| Osteopontin/OPN | Cell migration, adhesion and survival |
| VEGF | Encourages formation of blood vessels |
| G-CSF | Stimulates granulocyte and stem cell production and release |

**Supplemental Table 4.** PDL and doublings per day values of each cell line at each passage.

| Cell Line/Passage | Population Doubling Level (PDL) | Doublings per Day |
| --- | --- | --- |
| iMSC P1 | 5.7 | 0.63 |
| iMSC P2 | 11.7 | 0.74 |
| iMSC P3 | 17.1 | 0.68 |
| BM71 P1 | 3.9 | 0.56 |
| BM71 P2 | 8.6 | 0.67 |
| BM71 P3 | 12.7 | 0.59 |
| BM182 P1 | 4.1 | 0.59 |
| BM182 P2 | 7.9 | 0.47 |
| BM182 P3 | 11.5 | 0.45 |

**Supplemental Table 5.**
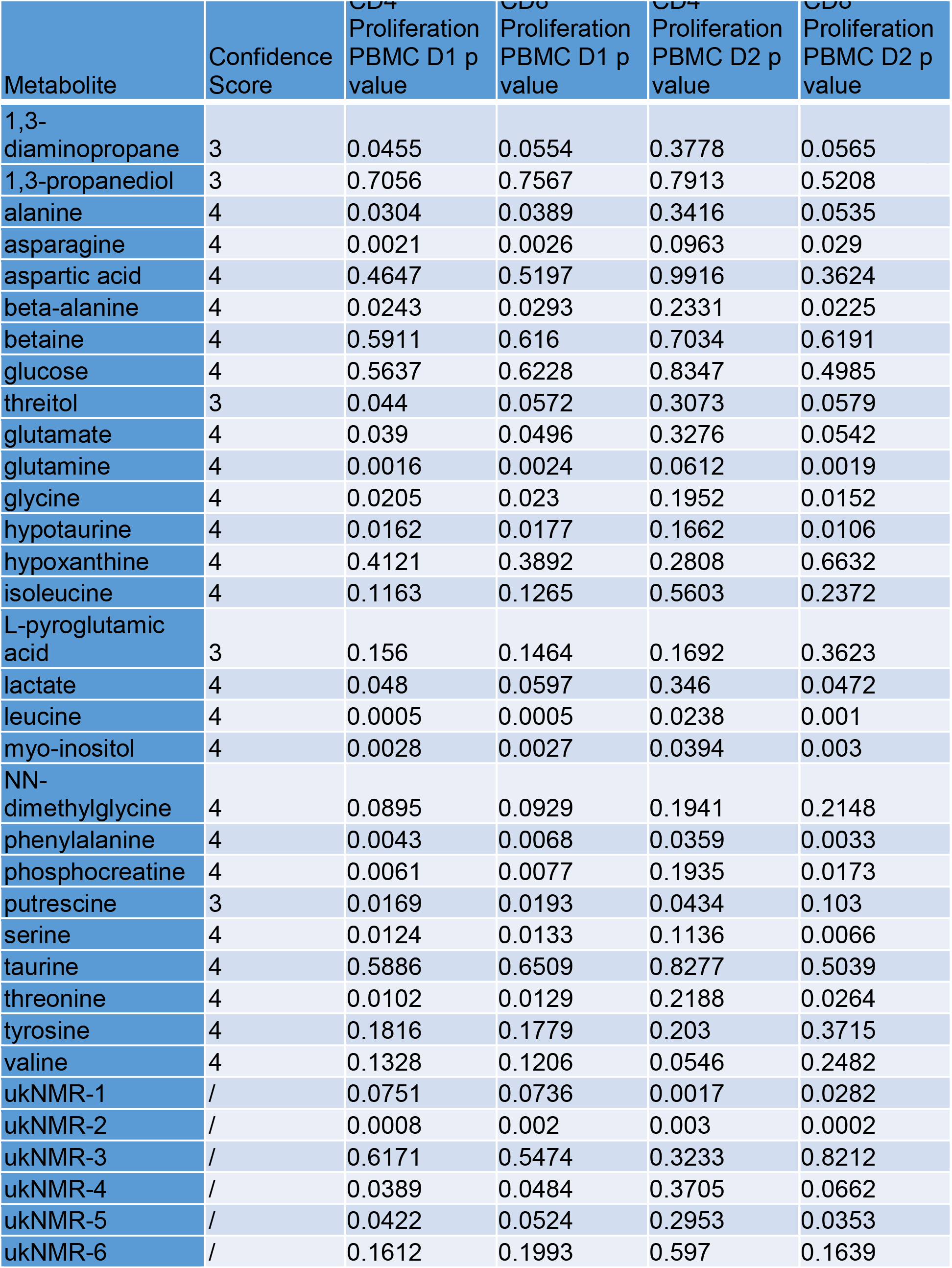
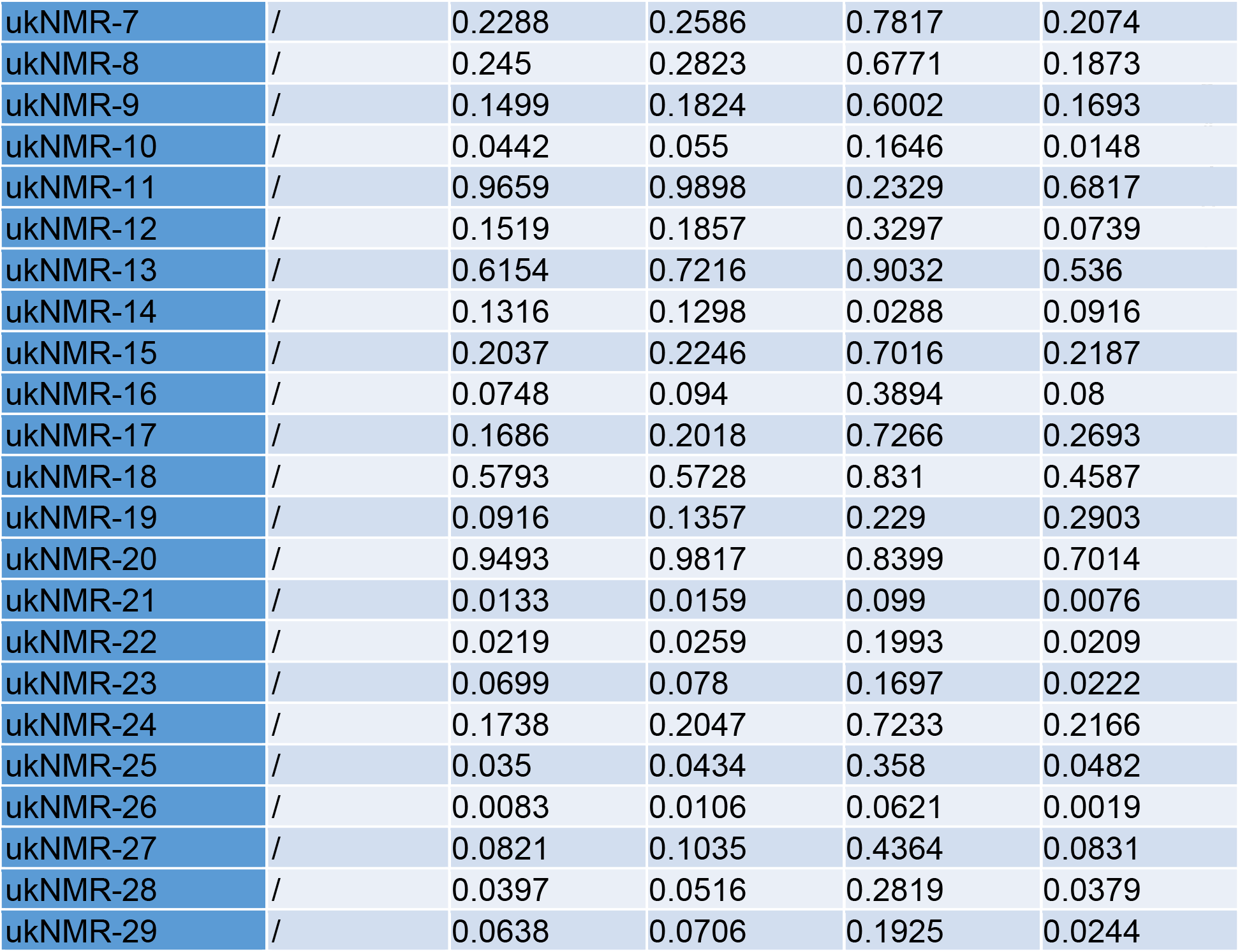
NMR identified metabolites, confidence scores and p-values from linear regression model. The metabolites were assigned a confidence level ranging from 1 to 5 according to published criteria.^32^

**Supplemental Figure 3.**
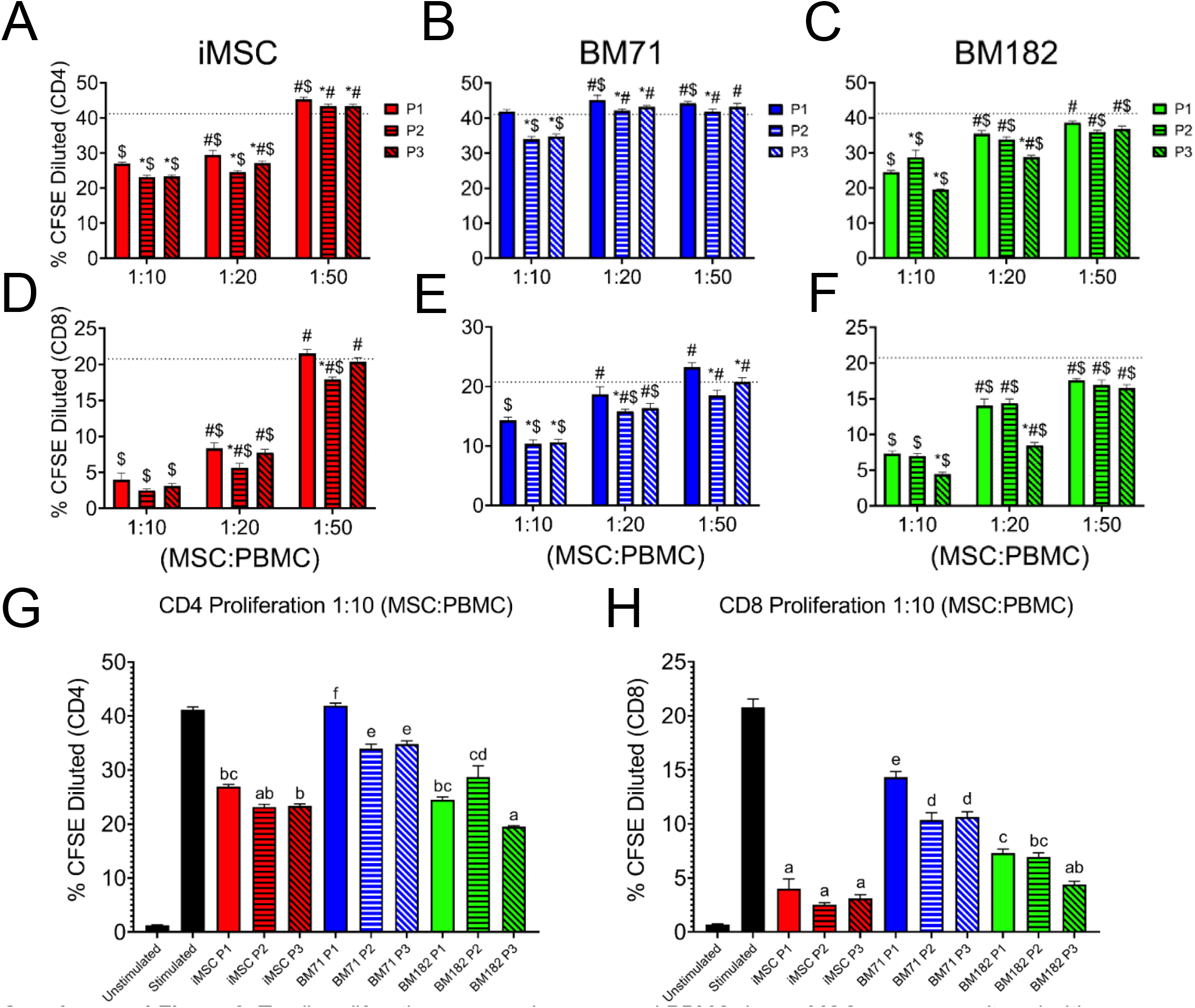
T cell proliferation assay using a second PBMC donor. MSCs were co-cultured with stimulated PBMCs at three different MSC:PBMC ratios (1:10, 1:20, 1:50). (A-C) CD4^+^ and (D-F) CD8^+^ T cell proliferation was assessed at each passage and ratio by % CFSE dilution. A 2-way ANOVA was used in order to determine if there was a significant difference from P1 within a ratio (*), a significant difference from the 1:10 ratio within a passage (#), or a significant difference from the stimulated control (dotted line)($) (P<0.05). (G) CD4^+^ and (H) CD8^+^ T cell proliferation comparing all cell lines and passages at the 1:10 ratio

**Supplemental Figure 4.**
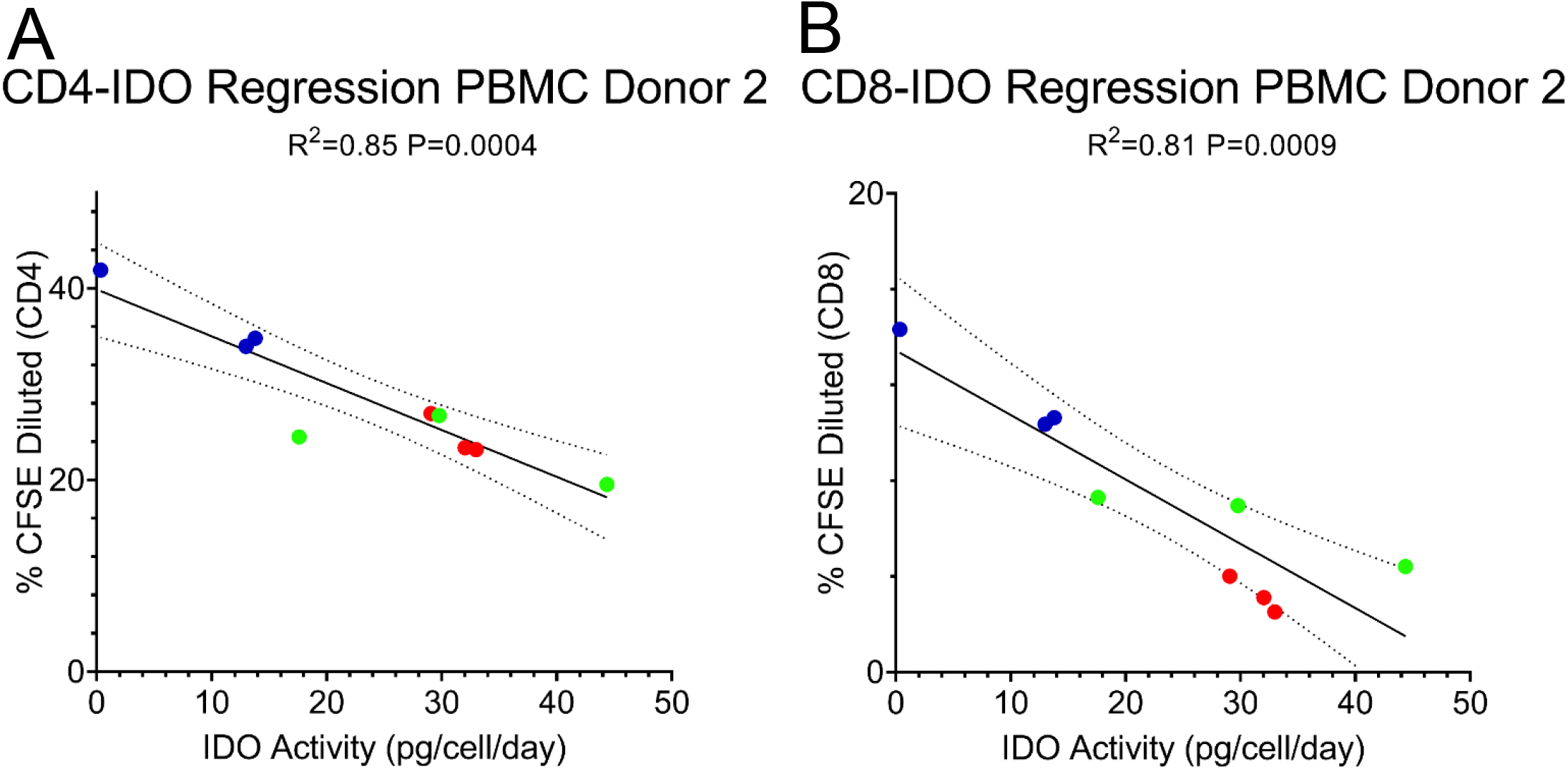
Regression of IDO activity and T cell proliferation with a second PBMC donor. Linear regression of the relationship between IDO activity and (A) CD4^+^ and (B) CD8^+^ T cell proliferation.

**Supplemental Figure 5.**
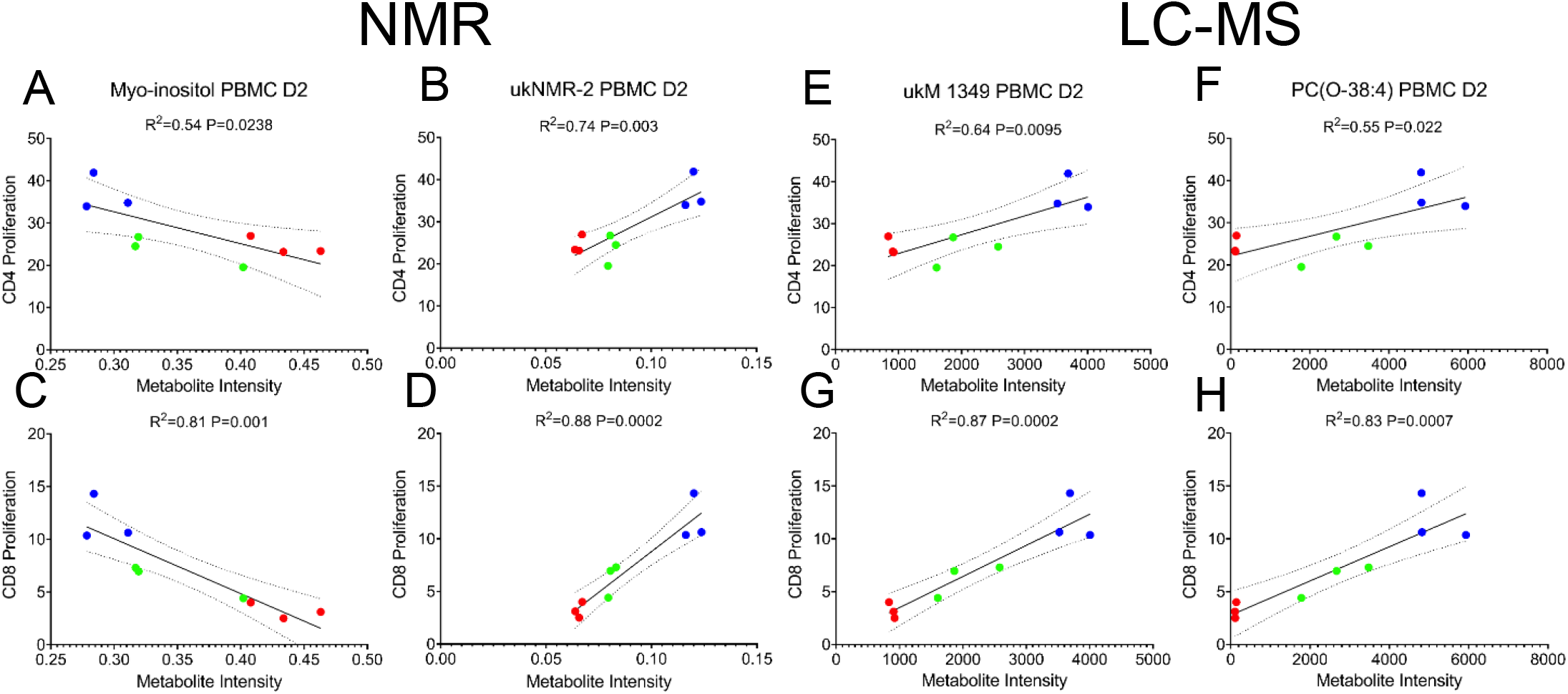
Linear regression analysis of the top correlated metabolites with PBMC donor 2. (A,C) Myoinositol, (B,D) ukNMR-2, (E,G) ukM-1349, and (F,H) PC(O-38:4) all correlated with CD4^+^ and CD8^+^ T cell proliferation for both PBMC donors.

